# 1/ A technical semi-field methodology to measure the effect of nutrition on honeybee brood rearing

**DOI:** 10.1101/2025.07.01.662504

**Authors:** Rui F.S. Gonçalves, Raquel T. de Sousa, Daniel Stabler, David M.S. Pinto, Geraldine A. Wright, Sharoni Shafir

## Abstract

A honeybee colony’s well-being is its ability to nurture larvae into healthy adults. Understanding how nutrition supports brood rearing is crucial for developing diets that could aid against environmental threats. Nutritional research on whole colony brood development has been historically challenging due to difficulties documenting the diet’s impact on brood production over time. We describe a novel semi-field method to study the influence of nutrition on brood rearing using standardised small colonies formed *de novo* (*ca.* 1500 nurse-age bees and a queen) housed in adapted mating-nucs, placed inside an enclosure and limited to feeding on chemically defined diets. Complete assessments were conducted every fifteen days, assisted by a bespoke device to photograph every frame to measure cell contents. A novel metric describes the number of bees generated per gram of diet consumed, measuring the impact of nutrition on brood rearing and overall colony size.

## INTRODUCTION

The honey bee colony’s capacity to produce brood and generate healthy young workers is a hallmark of robust colony health ^1–3^. The reproduction of honeybees exhibits a haplodiploid eusocial structure where adult individuals (male, reproductive female and unreproductive females) play distinctive roles in colony survival ^4,5^. Beyond the queen’s polyandrous mating, the population’s survival relies on the workers ability to maintain larvae, collect food, thermoregulate, and fight off pathogens and pests through their specific division of labour ^6,7^. Foragers collect pollen and nectar from flowering plants, which are processed by nest bees and stored as bee bread or honey, respectively ^8–10^. Nurse bees with engorged hypopharyngeal glands, between 3-12 days old, metabolise food to feed jelly to larvae and the queen ^9,11–13^. And colony workers regulate the colony’s temperature around 35°C for proper brood development ^14,15^. These tasks require adequate nutrition and precise coordination between nestmates ^16^. However, anthropogenic drivers have altered the diversity and abundance of flowering plants in their foraging landscapes, thereby affecting the health of honeybee colonies ^17–20^. Starvation conditions are becoming increasingly common, motivating beekeepers and scientists to develop feeds that mimic the composition of floral pollen to feed bees ^21–23^. Providing sucrose syrup in the absence of nectar flow, along with protein patties to promote brood rearing ^21,24^, has raised beekeepers’ operational costs related to feeds from 1% in 1976 to 20% in 2018 ^25^.

Research on colony nutrition has been ongoing since the 1930s ^26^, covering a variety of methods in settings such as open fields ^22,27–34^, semi-fields ^8,23,26,35–40^, labs ^41–43^, and complex indoor bee-flight rooms ^44–46^. Among the three primary macronutrients (carbohydrates, fats, and proteins), proteins play a crucial role in promoting brood-rearing ^21^, and pollen, the most studied protein source for honey bee colonies, has a substantial positive impact ^47^. However, certain pollen types, such as dandelion and maise ^35,40^, are insufficient to support colony growth, and a diverse pollen source is widely regarded as essential for maintaining healthy bee populations ^48,49^. When protein sources like pollen are restricted, brood-rearing ceases^50^, and if carbohydrate stores, like honey, are absent, the colony enters a period of starvation, putting its survival at stake^19,21,51–53^. Without pollen, colonies cannot raise brood.

The unreliability of pollen access has led to the development of diets rich in protein and other essential nutrients, which are often absent from natural pollen – referred to as pollen substitutes. However, these still do not appear to demonstrate a clear benefit to colony health^33^. Pollen substitutes are less effective than those diets containing at least 10% natural pollen in sustaining colony populations over the long term ^22,33,54,55^. Furthermore, when feeding these diets under conditions of pollen scarcity, bees deplete their internal reserves to support larval development, as they often lack essential nutrients, such as sterols ^21,24,52,56^. It remains unclear how long a pollen-free diet can support healthy populations throughout their annual cycle.

In 22 field studies on pollen substitutes, only five showed increased brood production ^57^. While valuable for understanding real-world conditions, field studies come with significant challenges. They are prone to uncontrollable variables (environmental, technical, or genetic) that affect brood-rearing more than feeding protein diets ^32,58^. For example, access to flowering plants and their pollen, even in small quantities, creates a confounding factor in brood rearing, which masks the nutritional effects of the tested diets ^54^. Methods to restrict access to foraging pollen, such as pollen traps ^33^ harm colony health by damaging workers^59^. Hive location may also expose them to plant protection chemicals, affecting colony development ^60,61^. Likewise, using large colonies can lead to the accumulation of other toxins^62^ used to control varroa mites, affecting capping rates, larval survival, and bee mortality ^63,64^. They are also more challenging to control for comb age; new combs have larger brood areas and heavier worker bees than older ones ^65^. Even the spatial distribution of bee bread within the comb affects brood-rearing success ^66^.

In semi-field studies, pollen access and foraging are effectively controlled. However, most use fewer than three colony replicates and rarely track brood over time. Colonies are typically confined in small outdoor cages (< 70 m³) ^67,68^, which increases mortality and reduces brood rearing ^36^. Colony sizes vary from 2,000 ^40^ to 10,000 bees ^68^ and the variety of methods for brood counting, such as visual estimation ^69^, sampling a central area ^70^, or image analysis tools ^71^, offers different levels of precision in estimating the diet’s impact on brood rearing.

Most importantly, comparing the nutritional impacts on brood rearing between studies is challenging without standardising methods and standard measurable metrics. In contrast, standard metrics are employed in honey bee toxicology to evaluate brood rearing. For example, the Oomen bee brood feeding test ^29^ and the OECD75 guideline ^72^ are widely used by industry and academic groups that test diets laced with toxicants, measuring the impact of pesticides on brood – via the brood termination rate metric (BTR), which measures the failure of eggs to transition into adults ^72^. An incidental outcome of these studies was the recommendation to use large polytunnels (>100 m³) to mitigate the harmful ‘caging effect’ on colonies ^66,73–75^. While BTR focuses on brood toxicity, comparable standards are needed to measure the positive impact in diet brood-rearing studies.

Therefore, we developed a technical semi-field method to address this need for standardisation and introduced a key metric, ‘bees per gram’, which measures the number of bees produced per gram of diet consumed. This metric enables the comparison of the brood-rearing potential of different diets across studies, locations, and time, in free-flying colonies under completely pollen-restricted conditions. Our experimental set-up used mini-colonies with 1,500 mixed nurse bees, five mini-frames with combs built the same year (without bee bread), and a same-year mated queen. Each mini-colony was housed in repurposed polystyrene mininucs, with 0.002 m³ of nest volume and a feeding chamber. And enclosed within a large flight arena – a polytunnel with 628 m³ of flight space, or within a 125 m^3^ glasshouse. Sucrose solution feeders were dispersed throughout to simulate natural nectar foraging behaviour. At the 15-day experimental mark, the brood assessment day utilised a custom device to photograph frames and analyse cell contents (empty, honey, or brood). This approach minimised confounding factors (e.g., pollen access, colony size, queen or comb age, foraging, and agrochemicals), allowing for a precise evaluation of diet effects on brood rearing.

In the present study, we developed an optimised brood assessment framework, showcasing key techniques that minimise disturbance to the colony while providing precise data on brood development. Our practical, reproducible approach streamlines the transition from laboratory nutritional studies to practical applications that can directly benefit beekeepers. We hope it serves as a foundation for standardising future whole-colony nutrition research.

## MATERIALS AND METHODS

### Nutritional bioassay

Our nutritional bioassay is based on queen-right honey bee colonies exclusively consuming artificial diets, which are then metabolised by nurse bees to feed their worker larvae. The developmental cycle of workers, from egg until emergence, usually takes about 21 days. Our assessments track this cycle almost to its end, from egg until capped pupae - three days before emergence (Fig. 1A).

**Fig. 1.**
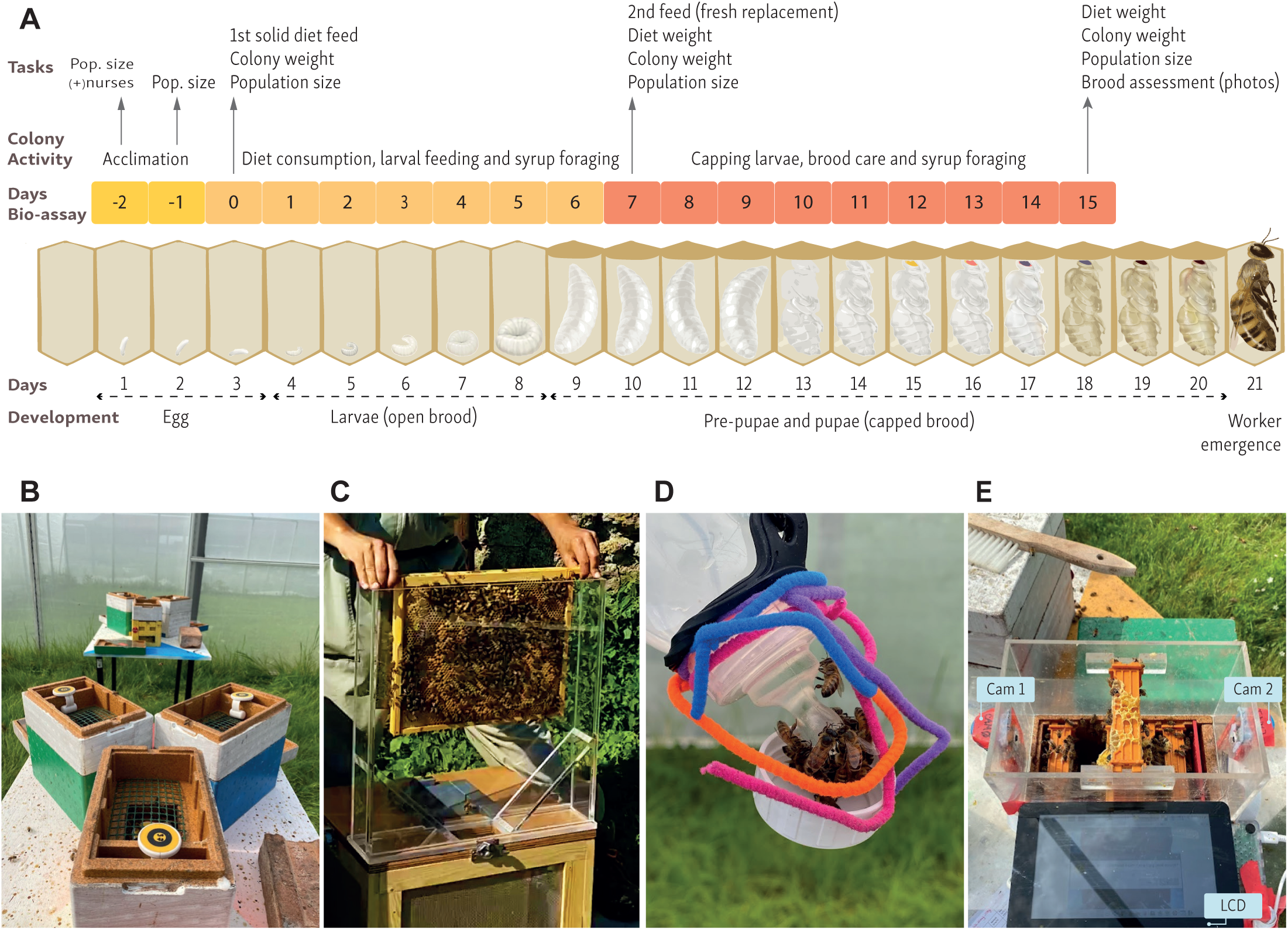
The brood assessment bioassay. **(A)** The chart matrix displays the honey bee development stage each day, stacked with bioassay days. Negative days represent the acclimation period, and days 0-15 represent the feeding period, which occurs within a 15-day cycle, followed by a final brood assessment. From left to right, the main tasks are: At day 0, assessing colony weight, population size, and feeding each with a solid diet (patty). On day 7, the colony weights, population size, and the weight change in the patty are used to estimate diet consumption. On day 15 (one brood cycle), we assessed the same metrics, plus a full brood observation, by photographing every frame across all treatments for a capped brood count. It also describes the mini-colony’s main activity: after acclimation inside the semi-field enclosure, it progressively consumes the diet to feed the larva, forages for nectar syrup, and ultimately takes care of its brood. All treatments started at the same brood stage – egg. **(B)** Mini-colonies in groups of three nucs in a triangle-shaped configuration; the picture displays the solid food chamber with the mesh supporting diet and a temperature sensor. **(C)** Nurse bees are collected and placed into the “nurse collection box” to replenish each mini-colony before the experiment begins. **(D)** Honey bees forage on artificial flowers with attached coloured velvet wires and bottle caps for support. **(E)** A novel, custom-made device built from acrylic sheets is designed as a box stand to support a Raspberry Pi LCD, two cameras, and displays a bee frame ready for photography.

Our method comprises three parts: acclimation, feeding and brood assessment. An initial set-up phase, represented in Fig. S3, is required to first establish the mini-colonies inside the controlled flight enclosures. This initial set-up takes 5 days, where day 0 is the beginning of feeding. The feeding cycles repeat from days 0 to 15. Consumption and the bee population are monitored during the 15-day cycle, and a full brood assessment is conducted at the end. If, in between the 15-day feeding cycles, the mini-colony consumes the diet and has more than two frames of bees, the system progresses through the feeding cycle uninterrupted. However, if the bee population criterion is unmet, it triggers a nurse addition stage and colony acclimation. This is an additional 2-day process that involves adding nurse bees to low-populated mini-colonies, and the system resumes the standard 15-day cycle. This dynamic ensures that the system remains adaptive and maintains a colony baseline throughout its cycle. The method operates in a continuous loop with a repeating 15-day feeding cycle; however, we conducted only one cycle in this study.

### PART I - Laboratory diet design

Our chemically defined diets consisted of protein, carbohydrates, fats, sterols, micronutrients, and a maltodextrin filler (with no nutritional value for the bee). We used as a protein source a plant-based - Soy Protein Isolate (Myprotein, UK; 90% purity) product as described in Sereia et al. 2013^76^. Here, we selected six diet treatments varying only in protein content (P): 6% (P6), 12% (P12), 15% (P15), 18% (P18), 25% (P25) and 30% w/w (P30). The percentage (w/w) of the diet total carbohydrate mixture ranged from 38% on P30 to 68% on P6, depending on the diet treatment. The carbohydrates consisted of about two-thirds of a mixture of 80% w/w sugar solution (in an equal mix of glucose and fructose) and one-third sucrose powder (Kent Foods, UK). As the fat source, we used soy lecithin (Optima Lecithin Health & Nutrition, UK) at 6% w/w of the total diet on all treatments as in *Stabler et al, 2021*^77^. We used Cholesterol (Sigma-Aldrich, ≥99%) at 0.1% w/w of the total diet as the sterol source, adapted from Herbert and Svoboda’s 1980 diets^78,79^ We then mixed all ingredients into a dough and kneaded them into 50-60 g (5 mm thick) patties. To test the pliability and texture of the dough formulations (structure tests), we further tested them by placing samples on top of a plastic mesh (lz 1 cm) inside an incubator at 34°C and 60% humidity (simulating in-hive conditions). After 24h, we discarded the diet formulations that absorbed excessive moisture and turned into sludge. We finally selected diets that successfully passed the structure test and that had been previously tested in our lab for palatability, consumption, and bee survival (> over 90%) in ventilated acrylic boxes (unpublished data), monitored over a week as in de Sousa et al. (2022)^80^.

### PART II - Mini-colony preparation and set-up

We assembled 36 mini-nucleus styrofoam boxes with dimensions 24 x 15 x 16.5 cm (Apidea Vertriebs AG, Switzerland), fitted with three plastic mini-frames with an 11 x 11 cm new wax foundation and one internal liquid feeder with clay pebbles inside to prevent bees from drowning. Each mini-nuc (mini-nucleus box) was number tagged and painted in separate colours to facilitate homing behaviour ^81^. To measure enclosed temperature fluctuations, we added a temperature sensor data logger (Omega, OM-EL-USB-1) to the food chamber of a small subset of mini-colonies.

To minimise the influence of pathogens and viral diseases on our experimental nurse bees, our apiary colonies had been treated with oxalic acid and amitraz between February and April, and Varroa levels were monitored using the sticky board method. We then selected healthy colonies from our apiary that were free of visible disease symptoms (i.e. the presence of Deformed Wing Virus) and with no detectable Varroa infestation.On day -5 we started collecting a selection of mixed nurse bees from these colonies by gently shaking open-brood frames covered with nurse bees into our custom-made “wooden nurse collection box” (37.4 x 42.2 x 31.4 height cm) (Fig.1C). This box covered by insect metal mesh, had three internal frames, spaced at 6 cm one from the other, allowing bees to suspend in cluster formation to dissipate heat in the field. A detachable acrylic funnel sits on top, facilitating the brushing of nurse bees into it. Nurse bees were brushed while lightly sprinkling with a 33% sugar solution laced with a couple of drops of lemongrass oil (Neal’s Yard, London) to prevent them from flying and to encourage nestmate formation ^82^. Nurse bees must be brushed at a rate of 1.5 frames of open brood per mini-nuc. Once complete, the box rests for one to three hours inside a dark room at 10 °C to promote cluster formation. At the time of nurse collection, no Varroa mites were detected, and no signs of deformed wing virus or other disease symptoms were observed. Since colonies showing visible signs of pests or diseases are behaviourally compromised and may influence brood rearing, any bees or colonies carrying mites or exhibiting visible signs of DWV were excluded.

Next, we turned each mini-nuc bottom up, sliding open the floor panel and exposing the three plastic mini-frames with the foundation. Lining all mini-nucs (Apideas)in a row, we sequentially placed on each a caged mated queen with candy (Candito per Api, PIDA, Italy) between the 2^nd^ and 3^rd^ frames. Our mated queens were sister queens supplied by the same Buckfast breeder. Next we used a 500 mL glass beaker to shave the bees off the collection box frames, filled each mini-nuc with 450 mL of mixed nurse bees (approx. 1,018) inside, and then slid closed the floor panel. Once all mini-colonies were formed *de novo* with nurse bees and a mated queen, we fed 350 mL of 1:1 (water to sugar) syrup into the internal feeder. We placed all mini-colonies housed in mini-nucs inside a ventilated dark room at 10°C for 48h for further nestmate and colony formation.

### PART III - Mini-colony acclimatisation to semi-field enclosures

We conducted two experiments, the first in a glasshouse with 30 colonies and the second in a polytunnel with 6 colonies. After the 48 hours of nestmate formation (day -3), at sunset, we released 30 mini-colonies inside the two-room glasshouse (l: 10 m x w: 5 m x h: 2.5 m) with 125 m^3^ of total flight volume to acclimate to the semi-field setting and to test our five increasing stepwise protein diets (P6, P12, P18, P25, P30). The repurposed glasshouse consisted of glass panels with a galvanised steel frame structure. Each of the two rooms of the glasshouse had a long central table where we placed 15 mini-colonies in groups of three (triangle shape) with entrances pointing in different directions. Due to the inherent heating capacity of the glasshouse infrastructure and to promote air circulation, we installed a mesh shade covering the whole area of the glass ceiling, two commercial extraction fans on both ends of the glasshouse, three portable aircon units, and three oscillating pedestal fans.

The second experiment was conducted in a purpose-built polytunnel (NP Structures, UK) measuring 18.4 m x 8.23 m x 4.15 m, providing six times more flight volume (628.4 m³) than the existing glasshouse. The polytunnel was covered in white insect mesh, and we repeated the same method with the (P15) diet using 6 new mini-colonies, which were placed in different directions in two groups of three, each on two tables, as described above.

After 24h of releasing the mini-colonies inside the semi-field enclosures (day -2, see Fig. 1A), we expanded their nest volume by removing the internal liquid feeder, adding two frames with empty built-in wax comb, and a top solid food chamber. During this procedure, we checked for queen release and visually confirmed that all queens had been naturally released, after which all empty queen cages were removed. If, for instance, queens remained caged, we manually removed the candy plug and reinserted the cage to allow a slower, controlled release. Mated queens usually start laying eggs after 24h of their release. Additionally, we kept in stock empty frames with built-in combs to standardise for frame composition, replacing any of the initial three foundation frames that were not yet fully drawn. We then top-up with an extra 250 mL of mixed nurse bees (approximately 566 bees) from our “nurse collection box” (Fig.1C). The solid food chamber, used for top-up nurse bees and placing our protein diets, is made from an APIDEA super feeder (700 mL), into which we cut a central hole (10 x 8 cm) and covered it with a green plastic mesh (12.5 x 9.5 cm, lz1 cm) (Fig.1B). At this stage, a total of 700 millilitres of mixed-aged bees (corresponding to approximately 200 grams) and around 1,580 bees, were added to each colony, close to 5.2 bee seams. Each bee seam was estimated to have approximately 300 bees. This estimation is based on an average of 787 bees per 100 grams and 226 bees per 100 millilitres, calculated from 28 research papers, including our own measurements.

Next, we suspended two artificial flowers per mini-colony throughout the enclosures, made from adapted 250 mL infant feeding bottles by piercing five lz0.5 mm holes in its nipples and adding colourful velvet pipe cleaners for bee identification and grip (Fig.1 D). Each bottle was filled with 250 mL of 33% sucrose solution and hung upside down, functioning as a gravity feeder to promote nectar foraging. Our artificial flowers restricted bees’ drinking access to only five at a time, preventing hoarding behaviour. The sucrose concentration in our feeders is in the mid-range of nectar of flowers typically visited by honey bees ^83^, and extends foraging activity within the enclosures relative to using higher sucrose concentrations. Additionally, such syrup concentration maintains high internal humidity favourable to brood development in summer and engages nest bees in activities of nectar-to-honey conversion. Drawbacks of low syrup concentration were noted during periods of continuous rain and cold within the polytunnel, leading to an outbreak of chalkbrood, consistent with findings by Flores et al ^84^. We washed and replenished feeding bottles per week and as needed. If, for instance, bees hoarded syrup, completely filling most of the comb cells in the mini-nucs, thereby obstructing egg laying, we replaced those frames during our inspection days with our in-house stock of empty built combs. We also placed four tubs with fresh water and support rocks in each corner of the enclosures for water supply.

Our mini-colonies were considered ready when they consisted of a single 5-frame brood chamber with the additional top food chamber (Fig. 1B). Internally, colonies contained five frames of fully built combs populated with bees (approximately 1,500 individuals), sugar syrup reserves, and brood exclusively at the egg stage produced by a mated queen.

### PART IV - Mini-colony nutritional bioassay for brood assessment, colony weight and population size estimation

We assigned *n=*6 mini-colonies to one of each of five diet treatments (P6, P12, P18, P25, and P30) to the glasshouse enclosure and (P15) to the polytunnel enclosure. The assignment criteria of diet to mini-colony aimed to maintain a balanced population size distribution and entrance direction among treatments and reduce bias on sun exposure, orientation, and colony population. Before feeding the mini-colonies with the protein diets, we recorded the initial weight of each mini-colony with a field scale (Ohaus Scout Pro Portable). We also recorded the number of bee seams (0 to 6 bee seams - clusters of bees occupying the spaces between two combs, or comb and hive wall) as per *Chabert et al, 2021* ^85^ by photographing the colony from above to estimate the population size of each mini-colony. Each bee seam was estimated to represent approximately 300 bees to calculate the colony size.

For all treatments, we weighed each diet to 50-60 grams of dough, rolled them into 5 mm thick patties, and placed them inside the food chamber over the green mesh, which sits 1 cm above the brood frames. Bees could access the food from below and move within the food chamber (Fig.1B) to consume the diet. This marks the start of our nutritional bioassay experiment (day 0).

A week later (day 7), with minimal disturbance to the colony, we removed the patty and registered population size and diet consumption by tracking changes in the patty’s weight for all mini-colonies and diet treatments (Fig.1A), taking an observation note if bees were eating the patty. The treatments are discontinued if nurse-aged bees avoid the diet altogether, indicated by marginal changes in diet weight or ‘negative consumption’ – i.e. when patties are not eaten and instead gain weight due to moisture absorption. In both experiments, the feeding cycle proceeded since consumption was observed in all treatments. Therefore, in our study, a fresh patty was provided, and its weight was recorded before adding the new patty. The mini-colonies were weighed to estimate changes in sugar stores.

On day 15 of our bioassay experiment (one brood cycle), we performed a complete mini-colony assessment on all treatments. We assessed colony weight, population size, honey stores, diet consumption and brood stages on all frames (Fig. 1A). We measured the brood stages by photographing both sides of every frame using a bespoke photographic-acrylic device (Fig. S1 - supplementary). The device consisted of a custom-made acrylic box (internal dimensions: 22.3 x 14.5 x 13.2 cm, lz 4 mm of thickness) that could be mounted on top of an open mini-colony rim, with a one-frame stand at its centre. On each end, it displayed two RPi cameras (64MP Autofocus Camera, Arducam) focused on the frame’s centre, with both cameras connected to a Raspberry Pi 4 (designated as the imager). Another Raspberry Pi 3 (named “controller”) was connected to a touchscreen display (Official Raspberry Pi 7” Touchscreen Display) and to the “imager” via an Ethernet cable (Fig. 1E). The “controller” controlled the “imager” remotely via SSH using a shell script software “image-experiment.sh”. Detailed set-up instructions and the code used are available at github.com/carandraug/beehive-imaging under the GNU All-Permissive License. The device recorded the time taken for the complete assessment (five frames, 10 pictures per mini-colony serially numbered); an experienced researcher spent, on average, 5 minutes and 41 seconds per mini-colony from setting the device to removing it. We then visually counted the number of capped brood cells in each image using available software (Adobe Creative Cloud 2019-2024, version 6.4.0.361, PS -Count Tool). At this stage of our experiment, we measured the influence of nutrition during the “one-capped bioassay cycle” (Fig.1A). We defined each bioassay cycle as the period when all capped brood was originated from eggs that were laid under the experimental procedure. At the start of the feeding period, all treatments started at the same brood stage (eggs) at day 0, with no capped cells present.

Not all of the eggs that the queen lays will develop into adults. Nutritional stress may lead to oophagy, the consumption of eggs by nurse bees, and up to cannibalistic consumption of the young larvae ^21^. Developing brood that is sick may be identified by nurse bees, which will open capped cells and remove the sick bee ^86^. However, a healthy brood that reaches the capped stage will likely emerge as an adult. In the subsequent 15-day bioassay cycles, the previously capped brood will emerge, the newly laid eggs will turn into pupae, and a new cycle begins, guaranteeing that we are not double-counting the same capped cells or missing capped cycles. Hence, we can continue the experiment for additional bioassay cycles by sequentially repeating the procedures from Day 7 and Day 15, measuring each newly capped cell that is in the stage between the pre-pupae and the point before the pupae emerge. In this study, we terminated the experiment after the first bioassay cycle.

### Data, Statistics and Power Analysis

Statistical analyses of diet consumption, capped-brood-cell counts, protein consumption, and “bees per gram” were conducted in Python 3.12.4 (https://www.python.org) within VS Code, with assistance from GitHub Copilot. Models and related functions from Statsmodels 0.14.1 (https://www.statsmodels.org) with numpy 2.2.3 and pandas 2.2.3 were used for all tests. We removed three mini-colony replicates from the analyses that lost their queen and had zero capped brood at the end of the 15-day bioassay.

To analyse diet consumption during the nutritional bioassay, we measured the total diet consumed over the 15-day period. Protein ingestion was estimated by multiplying each diet’s protein percentage by its total consumption. We then calculated our metric, ‘bees per gram,’ by dividing the number of capped brood cells by the total diet consumed. This metric represents the number of bees reared per gram of diet that is consumed, based on the assumption that each capped cell will develop into a viable adult bee. A Pearson correlation matrix was used to assess relationships between the independent variable (diet type) and dependent variables, including capped brood, total diet consumption, protein ingestion, population size, and ‘bees per gram.’

Our main objective was to evaluate the effects of nutrition, specifically protein content, on brood development. We explored the relationship between protein percentage as a continuous variable with our response variables—consumption, capped brood cells, and ‘bees per gram’ by using the Ordinary Least Squares (OLS) regression model from the Statsmodels 0.14.1 package. We estimated the intercept of the regression line to make it more accurate for our biological data. Key outputs include: the coefficient (coef), which indicates the strength and direction of the protein influence; the p-value and the R² value, which quantifies how much of the variation in the response is explained by protein. We also used the OLS to perform a power analysis to estimate: i) the probability of accurately detecting the protein effect on the brood - power (1 - β) and ii) considering that in future studies, we require an 80% degree of confidence – what is the sample size needed. We used Cohen’s *f* formula √(*r*^2^/(1 - *r*^2^)) to calculate the effect size and set the alpha level at 0.05.

The relationship between bee population size and capped brood was analysed by averaging population size between the three time-points assessed during the feeding period: day 0, day 7 and day 15. Population trends were analysed using Graphpad prism 10.4.1. To analyse population trends throughout the days and population differences between diets, we employed its mixed-effects model with Restricted Maximum Likelihood (REML) estimation at a significance level of 0.05, assuming that the variance between treatments was equal. Specifically, the mixed-effects model evaluated population differences between diet treatments inside the glasshouse (P6, P12, P18, P25, P30) on each assessment day and assessed population changes over time for each diet. This analysis covered the feeding period (days 0, 7, and 15) and the acclimation period (days -2 and -1). We then used Tukey’s post-hoc multiple comparison test to determine population differences between days and treatments. Since population size in mini-colonies can be a confounding factor for rearing capped brood, and since we observed absconding events among mini-colonies inside the glasshouse, we set criteria to identify potential outliers. Specifically, if a mini-colony showed a significant difference in population size in our mixed-effects model between Day 0 and Day 15, and its capped brood measurement had a z-score exceeding the factor of 2, meaning a data point beyond two standard deviations from the mean, we have considered that measurement an outlier. The z-score (*z*) for each data point (*x*) was calculated using the formula: *z* = (*x* – *µ*)/σ. Where (*µ*) is the mean of the data and (σ) is the standard deviation. Those outliers were not accounted for in statistics but are represented in figures.

To analyse the relationship between the two enclosures, polytunnel versus glasshouse, we selected a subset of our glasshouse data, the P18 diet, to compare it to the diet that was tested inside the polytunnel, the P15 diet. Here, we used a simple t-test to compare these two groups’ means and establish if the difference between them is statistically significant. In this case, we used Graphpad prism 10.4.1 t-test model.

## RESULTS AND DISCUSSION

### Protein-rich diets increase both diet consumption and protein intake

The mini-colony’s access to pollen was entirely restricted and absent from all comb stores, with protein provided solely through the experimental diets delivered to the mini-nucs throughout the 15-day bioassay cycle. We performed an Ordinary Least Squares (OLS) model to evaluate the influence of protein percentage on diet consumption (Fig. 2A). The OLS showed that protein-rich diets positively affected diet consumption (β = 1.33, p < 0.005), indicating that for each protein percentage increase in our diets, our mini-colonies will eat 1.3 grams more. The model explained 35% of the variance in diet consumption (R^2^ = 0.35, F(1,25) = 13.6, p = 0.001.) Additionally, the Pearson correlation demonstrated a positive correlation between protein-rich diets and diet consumption (*r(25)* = 0.59, *p<0.005*). Although protein strongly influences diet consumption, it doesn’t necessarily mean colonies will neglect diets that are entirely absent of protein. For example, other dietary components, such as carbohydrates, can also significantly influence food collection and consumption in honey bee colonies^87–89^.

**Fig. 2.**
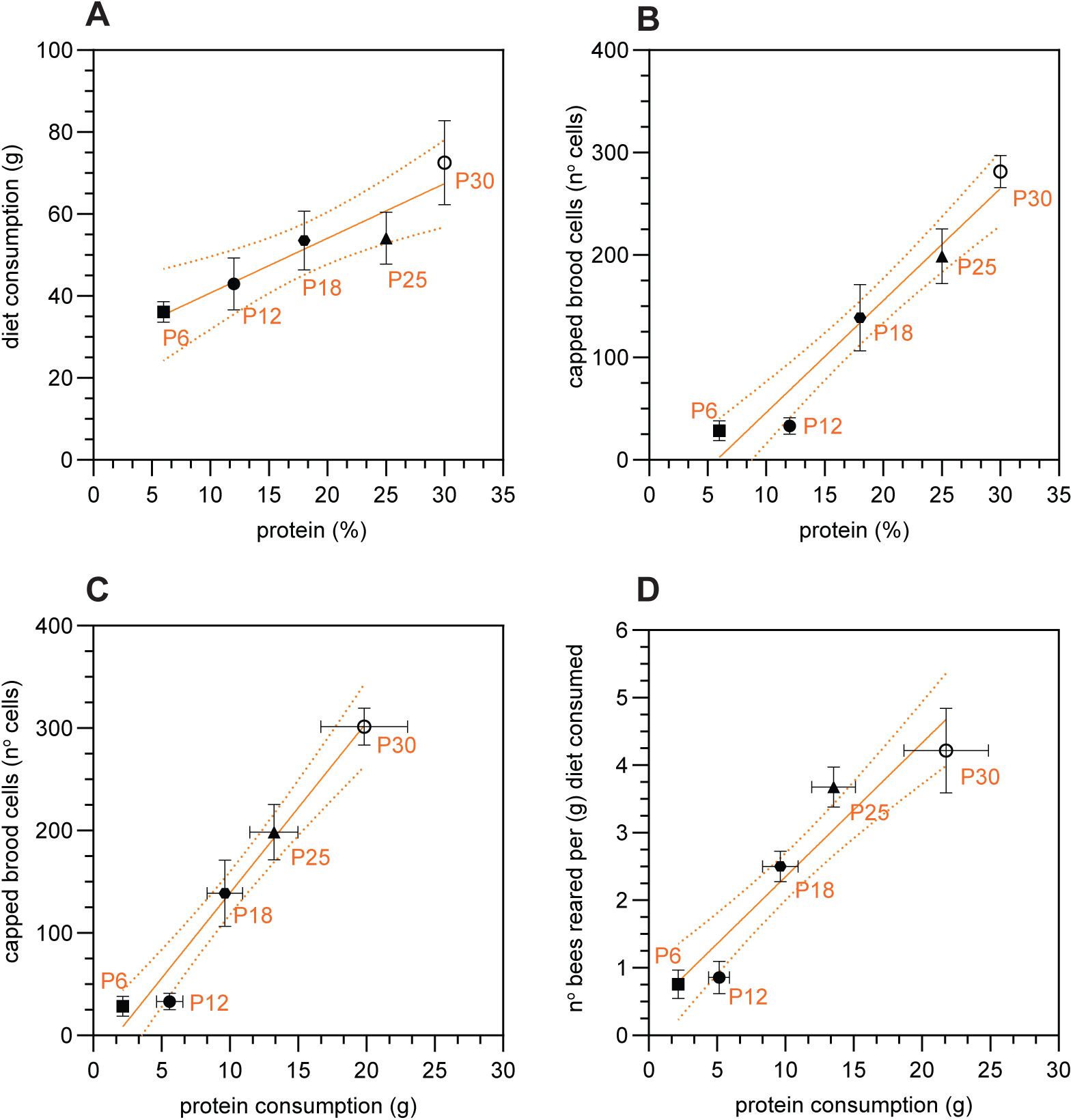
Protein content proportionally influences diet consumption, capped-brood rates and the number of bees reared in mini-colonies. Data points in all figures show the average results for each of the five diets varying in protein content (% w/w) inside the glasshouse: P6 (6%), P12 (12%), P18 (18%), P25 (25%), and P30 (30%) - over the 15-day period (one bioassay cycle). Orange lines depict the regression lines with 95% degree confidence between different diet types. Error bars in SEM, *n=5* mini-colonies per (P6), (P12), (P30) diet; and n*=6* mini-colonies (P18), (P25) diet. **(A)** The relationship between dietary protein percentage and total diet consumption over the 15-day period. A positive linear association was found, with the OLS model showing that higher protein content is associated with greater consumption. **(B)** Relationship between dietary protein percentage and the total number of capped brood cells counted at the end of the 15-day period. A positive association was found, showing that higher protein content significantly predicted more capped cells. **(C)** The relationship between protein intake (grams) and the number of capped-brood cells (count) shows a positive association with protein-rich diets, leading to higher protein intake and a higher number of capped cells. Our linear regression shows that for every gram of protein ingested, mini-colonies can generate 12 more bees **(D)** The relationship between protein intake and our metric “bees per gram” (number of bees reared per gram of diet consumed). Mini-colonies on protein-rich diets like P30 were, on average, able to rear 4 bees per gram of diet consumed, while lower protein-rich diets (P6 and P12) produced fewer than one bee per gram of food consumed.

Since diet consumption and protein intake are inherently linked in our experiment, mini-colonies that consumed more of the protein-rich diets also ingested more protein (Fig. S1). Here, the OLS model showed a strong fit, explaining 80% of the variance in protein intake (R² = 0.80, F(1,25) = 104.2, p < 0.001). The regression coefficient indicated that for every gram of diet consumed, colonies ingested approximately 0.35 g of protein (β = 0.35, p < 0.001). A Pearson correlation further confirmed a strong positive association between diet consumption and protein intake (r(25) = 0.90, p < 0.005) (Fig. S1). These results indicate that mini-colonies consumed more and ingested significantly more protein when offered protein-rich diets (Figs. 2A, S1), suggesting that dietary protein may have both phagostimulatory and metabolic effects that enhance consumption.

### Protein-rich diets strongly influence the number of capped brood cells and produce significantly more bees per each gram of diet consumed

Protein plays a critical role in the ability of a honey bee colony to produce healthy young workers as brood rearing ceases without an adequate protein source. One of our primary objectives was to assess how dietary protein content influences brood production and whether a stepwise increase in protein percentage drives greater protein intake and brood rearing. As shown in Figure 2B, we observed a clear positive relationship between dietary protein percentage and the number of capped brood cells. The OLS demonstrated a strong fit, explaining 76% of the variance in capped brood production (R² = 0.76, F(1,25) = 78.61, p < 0.001). For every 1% increase in dietary protein, colonies produced an average of 11 additional capped cells (β = 10.9, p < 0.001). A power analysis based on our sample size of 27 mini-colonies indicated a 99% probability (power = 0.99) of detecting a protein effect within the tested range. If in future studies a power of 80% were acceptable, only five colonies (N = 5), one per diet level, would be needed to detect a similar effect.

A consistent trend was observed for the relationship between protein intake and brood production (Fig. 2C). The OLS model showed a strong fit, explaining 74% of the variance in capped brood (R² = 0.74, F(1,25) = 69.56, p < 0.001). For every gram of protein consumed, mini-colonies produced an average of 12 additional capped brood cells (β = 12, p < 0.001). Protein-rich diets had a marked impact on brood production. Colonies fed the highest-protein diet (P30) produced more than twice as many capped brood cells as those on P18, and nearly ten times more than colonies on the lowest-protein diet (P6). These findings support our hypothesis that total protein intake—rather than overall diet consumption—is the primary driver of brood development. Diets with higher protein content significantly enhanced brood production, suggesting that increased protein availability improves colony productivity by increasing worker numbers, which may in turn benefit overall colony health.

The ability to precisely quantify both diet consumption and the number of capped brood cells enabled us to estimate diet efficiency, defined by our key metric: bees per gram—the number of bees produced per gram of food consumed (Fig. 2D). The mini-colonies fed diets with low protein content like (P6) and (P12) generated less than one bee per gram, whereas protein-rich diets like (P25) and (P30) reared four times more bees per gram of diet, 4.39±2.78 and 4.43±1.21 bees/g, respectively. This relationship between protein and brood is consistent with previous studies, such as Ricigliano et al. ^22^ and others ^30,34,90–94^, demonstrating a positive relationship between protein supplementation and brood development. Similarly, Li et al. (2012) ^95^ identified pollen substitutes containing 30-35% protein as the most influential factor for promoting bee development.

To our knowledge, this brood bioassay method is the first to estimate the number of bees a protein diet can produce and evaluate the influence of protein on brood development in a stepwise approach. The novelty lies in demonstrating that a stepwise increase in dietary protein content leads to increased diet consumption (Fig. 2A), and that greater consumption of protein-rich diets results in a compounded intake of total protein (Fig. S1). Furthermore, consumption of protein-rich diets was associated with a higher number of capped brood cells produced (Figures 2B and 2C), ultimately improving brood production efficiency, as evidenced by a greater number of bees produced per each gram of diet consumed (Fig. 2D). Future studies should incorporate a positive control using pollen-based diets and a negative control consisting solely of carbohydrates.

### Supplementation of nurse bees and population dynamics in mini-colonies: the population was affected by supplementation but not by diet treatments

Mini-colony populations were formed by mixing unrelated nurse bees into a single group, orphaning them by keeping them queenless for three hours and spraying them with sugar syrup laced with lemongrass ^96^. We measured population sizes during the acclimation period (days –2, –1, and 0) and standardised the colony population before feeding.

On day -2, at least 24 colonies required supplementation, with fewer than 1,500 bees, an average population size of ∼1,050 ± 146 bees. One day after supplementation (day -1), the populations increased to 1,530L±L202 bees (Fig. 3A; mixed-effects model, three time points; P < 0.0001). And on day 0, we observed a slight population reduction, likely due to disorientation and losses within the enclosure. Despite this, supplementation increased the population to a mean of 1,422 ± 195 bees on day 0, close to our target (Fig. 3B). This technique effectively standardised colony populations before the feeding period (day 0).

**Fig. 3.**
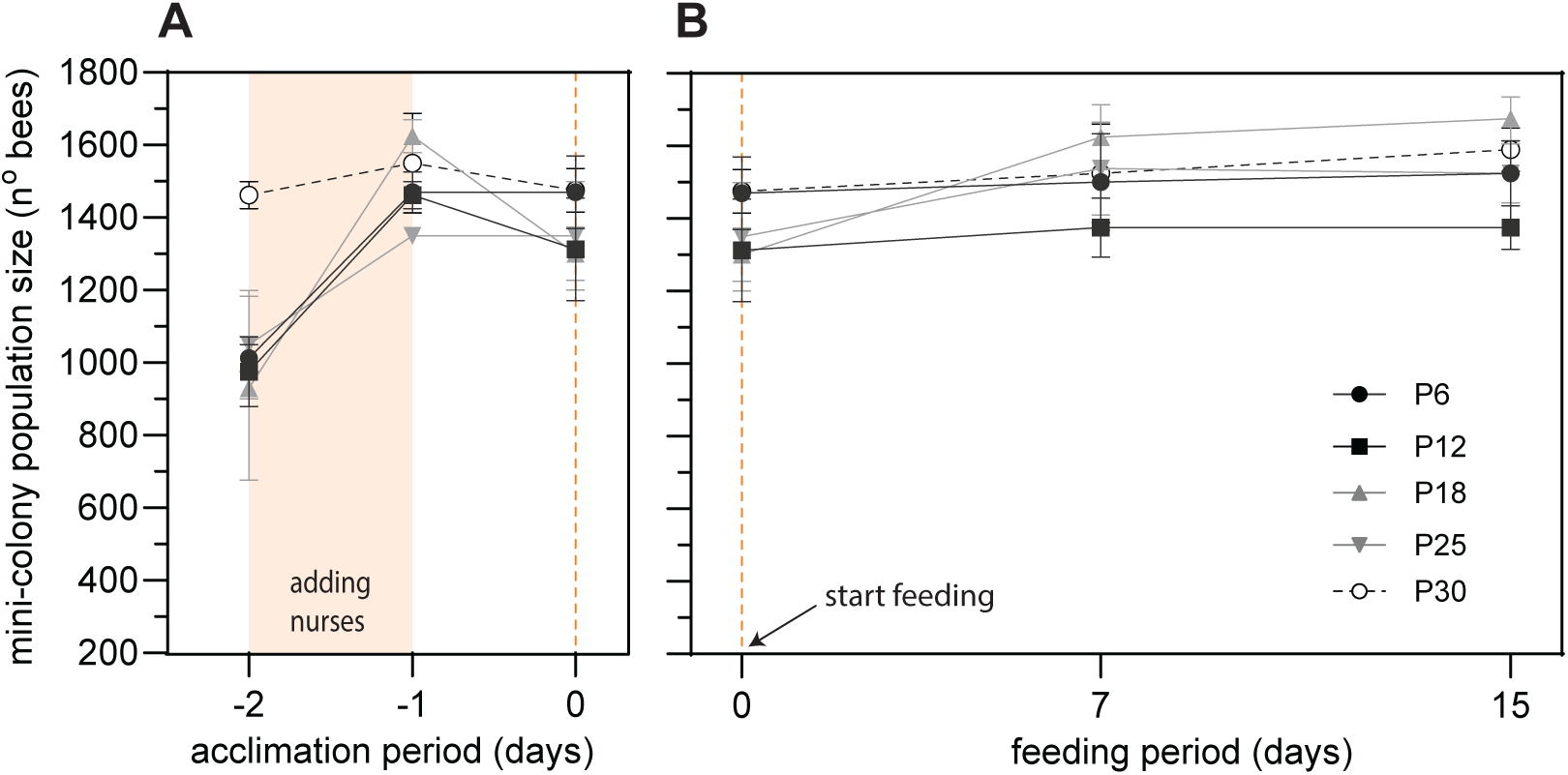
Population size remains stable during the feeding period. **(A)** Population dynamics inside the enclosure (glasshouse) during the acclimation period highlighting the addition of extra orphan nurse bees between day -2 and day -1 (shaded area) (∼500 bees) and start feeding day 0 (dashed orange line). Nurse addition increased the low populated colonies (Mixed-effects model (REML), Days *P*<0.0001), standardising colony populations between mini-colonies and diet treatments at the start of the protocol (day 0) with similar population sizes 1,400-1,500 bees (*ns*, Mixed-effects model (REML), Day 0 P>0.005). **(B)** Population size inside the glasshouse over the 15-day feeding cycle remained stable on most protein diets except on P18. There were statistical differences in population trends from Day 0 to Day 15, on the P18 diet (Mixed-effects model (REML), Days *P*<0.05, Tukey HSD; *P<*0.005) due to absconding bees randomly entering these colonies. There were no statistical differences between bee populations based on the diet type (*ns*, Mixed-effects model (REML), DIET *P*>0.05).

During the feeding period, between day 0 and day 15, we did not anticipate significant differences in population size over such a short timeframe unless elevated adult mortality were present. In fact, colony size did not differ significantly between diets (Fig. 3B; mixed-effects model, P > 0.05), and any influence of colony size on brood development in our semi-field mini-nucs is therefore likely minimal. Small colony size might still disrupt honey bee behavioural flexibility, and we cannot exclude the presence of precocious pollen foragers despite protein patties being available to each mini-nuc and natural flowers being absent in our enclosures. Our use of external slow-release artificial nectar flowers, rather than internal syrup feeders, aimed to maintain a functional equilibrium between foragers collecting nectar and housekeeping workers attending the brood.

Colonies were formed in two steps: on colony formation day, outside the enclosure with three mini-frames of bees, and on day -4 (24h after being released inside the enclosure) by adding two more frames and extra bees to expand the brood chamber. Previously, we attempted to form colonies fully on colony formation day with over 1,500 bees and five frames, but faced high mortality due to poor ventilation in the mini-nuc and limited syrup consumption during the critical 48-hour cold, dark period. To avoid these issues, we adopted the intermediate supplementation step on day -2. Regarding behavioural observations, orphaned nurse bees behaved like natural swarms, forming cohesive clusters with little flight and showing plasticity in nestmate acceptance. The “nurse collection box” effectively created and replenished mini-colonies. Brief aggression toward non-nestmate nurses was noted in a few colonies but resolved within two hours.

Assuming that the average lifespan of a worker honey bee is approximately 30 days, worker mortality will be more visible in these mini-nucs. In cases of extended bioassays, the periodic replenishment of mini-nucs, as described in the Methods, will help maintain a stable worker population, and because supplementation and absconding can affect colony nutrition, it is crucial to synchronise these events across treatments and in between the 15-day intervals, while also recording any instances of absconding. The pooling of nurse bees from multiple healthy donor colonies further minimises inter-colony variability in protein and lipid reserves, thereby standardising the overall nutritional status across treatments. Collectively, our results indicated that neither colony demography nor nurse nutritional variation significantly influenced brood-rearing.

### Enclosure systems strongly influence ambient temperature and significantly influence brood health

The method described was developed over five years, with trials involving different mating-nucs, enclosures, colony sizes, and feeder configurations, until it reached the refined stage presented here. Maintaining the daily average ambient temperature inside the enclosures below 37.5°C is crucial, as higher temperatures negatively affect brood survival^15^. For example, Xu et al. (2023) showed increased mortality in pre-pupae and pupae at 40°C ^97^.

In July 2019, the glasshouse ambient readings from the supporting tables (n=2) and in-hive temperatures (readings from the food chamber, n=2) were measured every 5 min during the hottest days (Fig.4). Peak ambient temperatures reached 45°C inside (Fig. 4A), leading to queen losses (n = 3). Despite the losses, the remaining mini-colonies were capable of regulating their internal temperature, maintaining a more stable and less variable environment, as demonstrated by the lower standard deviation (SD) between the inside (6.31°C) and the outside (10.32°C) temperature, as well as the smaller interquartile range (IQR); inside (11.5°C) compared to the outside (20.1°C). The misting system effectively mitigated some heat stress, activated in 30-minute bursts from 12:30 to 16:00, reducing the ambient temperature from 43 °C to 30 °C (Fig. 4A). Once it stopped, the temperature rose to 40 °C.

**Fig. 4.**
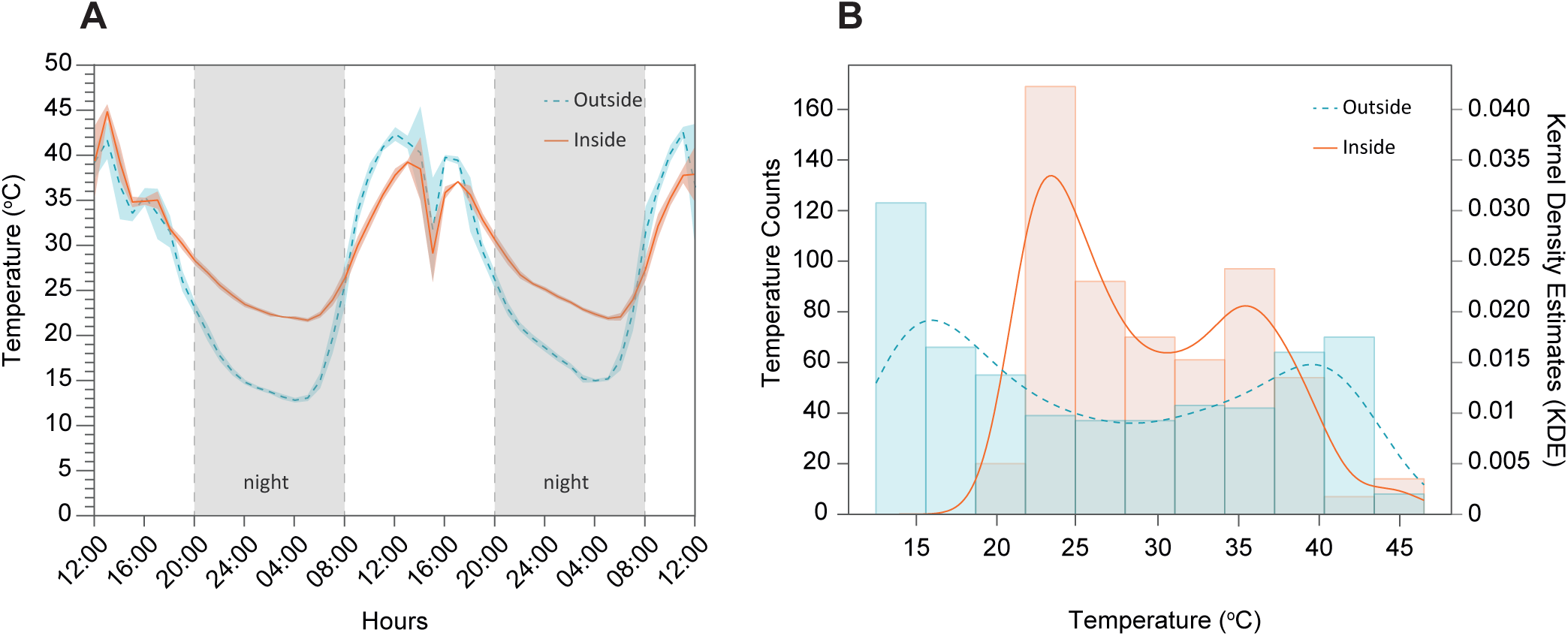
The mini-colony population was able to buffer environmental temperature fluctuations. In both figures, temperature fluctuations inside the mini-colony (orange) (*n=*2) and outside (blue) (*n=*2) over a 60-hour (2.5 days) period during a July heatwave inside the glasshouse. **(A)** Maximum environmental temperatures reached 45°C during daytime, and the minimum temperature was 14°C during nighttime (grey shaded area). The temperature drop at 16:00 resulted from the misting system’s action, which began at 11:00 and continued until 20:00 in 30-minute bursts, to cool the environment. **(B)** Histogram temperature variation between the inside (orange) and outside (blue), the columns represent the temperature histogram count chart (bin width =3), and the lines represent kernel density estimates (KDE), which are probabilities based on the histogram data. The hive’s internal temperature was more stable than the outside temperature, with a shorter range and less variability.

Different enclosures have different environmental conditions, significantly influencing the mini-colonies’ capacity to rear their brood as seen in Fig. 5A. Here, we excluded one replicate from the (P18) diet group, its capped brood had a z-score of 2.213, exceeding two standard deviations from the mean, and the mini-colony showed a significant population increase from Day 0 to Day 15. This met both of our criteria for identifying outliers. Yet, despite similar consumption and population sizes (Fig. 5B and 5C; unpaired t-test, ns, *P>*0.05), mini-colonies inside the polytunnel were able to rear twice as many capped brood cells compared to those inside the glasshouse (Fig. 5A; unpaired t-test, *P<*0.0001). The main methodological differences between locations were the type of enclosure and the protein content of the diet. As previously discussed, protein contents influenced brood-rearing capacity (Fig. 2B and 2D). And although the P15 diet in the polytunnel is lower in protein, it resulted in 59% more capped cells than in the glasshouse (Fig. 5A), likely due to the glasshouse’s higher temperatures (up to 40°C) and poorer ventilation.

**Fig. 5.**
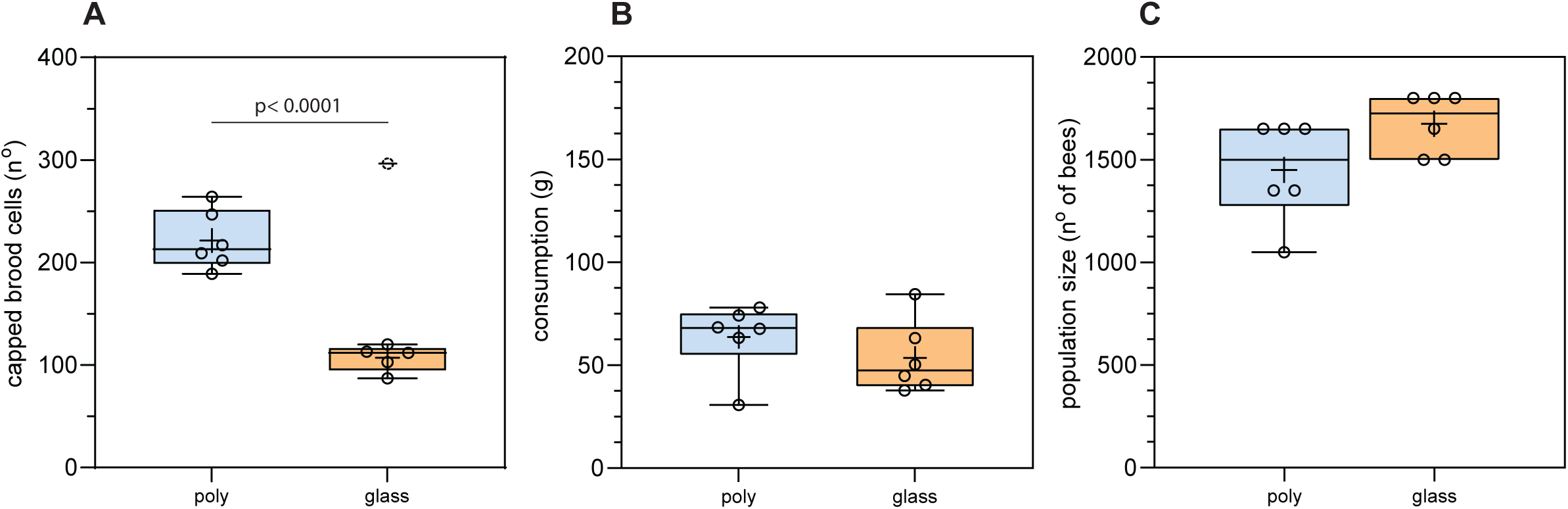
Semi-field conditions can influence capped brood production in mini-colonies. Comparison of mini-colony development inside the polytunnel (blue) and glasshouse (orange) during the first 15 days – one bioassay cycle. Colonies inside the polytunnel (blue) (n=6) were fed a diet with 15% protein (P15) (n=6), and inside the glasshouse (orange) were fed a diet with 18% protein (P18). **(A)** Mini-colonies in the polytunnel consumed diets with lower protein content compared to those in the glasshouse. The number of capped brood cells inside the polytunnel was two times higher than in P18 mini-colonies (unpaired t-test, p < *0.0001).* The dashed circle represents our flagged outlier, which was excluded from our statistical analysis. **(B)** Mini-colonies inside the polytunnel and the glasshouse showed similar diet consumption (ns), ∼60g during the first 15 days. **(C)** Mini-colony population size (ns) was also similar between both groups, with a mean difference of 225 bees.

### Considerations on temperature control, enclosure design and optimisation

Trials conducted inside the glasshouse proved significantly challenging in keeping temperatures below 37 °C. We have employed shading nets, misting systems, air conditioning units, pedestal fans, and two horizontal industrial extraction fans; however, these measures were insufficient to prevent some queen losses and absconding during heat waves. Additionally, the type of ground floor cover can impact the temperature and air quality within the enclosed space. Ground covers made of concrete slabs have high thermal mass, leading to the accumulation of bee carcasses, which increases the researcher’s susceptibility to heat stress and bee dust-related fever, necessitating regular cleaning. In the purpose-built polytunnel (628 m³), the ground was intentionally covered with perennial ryegrass, improving heat absorption and ground sanitation – bee carcasses naturally and quickly decomposed. Flowering was prevented by regular mechanical mowing of any appearance of flower buds. Furthermore, other differences likely affected colony well-being. The polytunnel had higher flight activity, better ventilation (netted walls), and a natural floor, unlike the glasshouse’s concrete slabs. We recommend using enclosures that more closely mimic natural environments, such as polytunnels, whenever possible.

Other observations made inside the enclosures were foragers’ ability to collect sugar syrup from various feeder types and deliver it to their colony. Feeders providing ample drinking space were emptied within hours, causing some mini-colonies to fill rapidly. When we supplied high-concentration syrup (>50% carbohydrates), we noticed a reduction in the area available for queens to lay eggs. In these mini-colonies, sugar storage can rapidly occur in the middle frames, limiting the number of empty cells available for egg laying and potential brood rearing. A syrup concentration of 33% carbohydrates, similar to the nectar found in wildflowers^98^, was preferred. However, when artificial flowers ran out of nectar, foragers attempted to expand their foraging range, with many repeatedly contacting the enclosure walls, which likely increased exhaustion and mortality. During this period, light diffusion through the enclosure caused some foragers to become trapped against the walls brightest areas, effectively forming “light traps.” Restricted foraging may alter colony dynamics; however, the enclosed system was designed to reduce environmental variability and allow for a focused test of nutritional effects on brood. This trade-off highlights the balance researchers must strike between experimental control and ecological realism when studying colony-level nutritional responses.

Future studies should explore methods to mitigate this behaviour by keeping foragers engaged in nectar foraging activities, improving enclosure design through enhanced light dispersion or shading, and implementing more effective cooling systems to mitigate heat shocks during the summer.

### Development and optimisation of a minimally disruptive image-capturing system for standardising honey bee nutritional studies

The need to shake colony frames to release adult bees for image capture, combined with sudden fluctuations in comb temperature, can result in high stress levels for both adults and brood, negatively impacting colony health. The novel device, specifically designed for this bioassay method, sits on top of the mini-colony, allowing us to optimise image timing and minimise disruptions to our mini-colonies during the 15-day comprehensive assessment cycle. Frames with adult bees are shaken sequentially into the inside of the acrylic box, guaranteeing that all nurse bees are placed back in their hive with frames to support them. The acrylic box is made with off-the-shelf anti-glare Perspex acrylic with UV protection, similar to what museums use to protect picture frames. This provides some UV protection to honey bee eggs and larvae, which are sensitive to UV irradiation ^99^.

The image capture speed on both sides of the frame is approximately ≈1.19 min per frame. A single experienced researcher can complete a full assessment of each mini-colony in (≈ 6.35 min), less than 10 min per mini-colony. Previously, two were needed to speed up the process and reduce colony disruption. To the best of our knowledge, this is the quickest and least disruptive method to capture images across all honey bee colony frames. The image’s size is approximately 2.2 MB, and its quality is assessed by visually confirming it on the touchscreen display. We repeated image capture if images turned out to be out of focus, stray bees covered the combs, or environmental light impeded a clear image (Fig. S2). We captured 800 frame images in one trial, and 39 of them were out of focus or poor quality. Counting cells with eggs and larvae remains challenging, as the device only captures the full depth of the comb located at the centre of the mini-colony frame, about 100 cells per frame side. This is due to the natural individual cell shape that has a hexagonal structure tilted upwards at 9-14° and with a depth of aprox. ≈12mm. In contrast, capped brood and honey cells are easily quantifiable by visually counting each comb cell surface.

The double Raspberry Pi set-up was required because the Raspberry Pi 4 cannot reliably and simultaneously control the Arducam 64MP Hawkeye cameras and the touchscreen display. Therefore, the two cameras were connected to one Raspberry Pi (RPi), the touchscreen display was connected to the second RPi, and an Ethernet cable connected both RPi. However, improvements are still possible in further iterations to increase capture speed and image quality. For example, installing additional cameras would allow us to capture comb edges more effectively, or a scanning method (with a single camera) would enable us to examine each frame from top to bottom. Additionally, making the system completely wireless provides higher flexibility and manoeuvrability in field-type settings. Future enhancements could also incorporate artificial intelligence to capture honey bee larvae more effectively and include small temperature sensors in the acrylic box to alert the researcher if local micro-temperatures that harm the brood are extremely low or high. We hope this method establishes a foundation for standardising future nutritional studies as a whole.

## Acknowledgements

The authors would like to thank Haim Kalev for his general beekeeping husbandry; John Hogg for crafting the nurse-collection and acrylic boxes; and Jennifer Chennels for laboratory support. This research was supported by grants from the BBSRC: BB/T015292/1, BB/P007449/1 to GAW and SS and BB/T004258/1, BB/R019614/1 to GAW. DMSP is supported by the EPSRC Programme Grant Visual AI EP/T028572/1

## Competing interests

The authors, Rui FS Gonçalves, Raquel T. de Sousa, Daniel Stabler, Geraldine A. Wright, and Sharoni Shafir, are shareholders in Apix Biosciences, a bee-nutrition university spin-off registered in Belgium. The authors declare that they have no other competing interests. David MS Pinto has no competing interests to declare. All authors have read and approved the manuscript for submission.

## Author contributions

**RFSG** (Rui Gonçalves), **RTDS** (Raquel T. de Sousa), **DS** (Daniel Stabler), **DMSP** (David M.S. Pinto), **GAW** (Geraldine A. Wright), and **SS** (Sharoni Shafir) contributed as follows: Conceptualisation—RFSG, RTDS, GAW, SS; Methodology—RFSG, RTDS, DS, DMSP, GAW, SS; Software—DMSP; Validation—RFSG, RTDS, GAW, SS; Formal analysis—RFSG, SS; Investigation—RFSG, RTDS, DS; Resources—GAW, SS; Data curation—RFSG, RTDS; Writing (original draft)—RFSG, SS; Writing (review & editing)—RFSG, RTDS, DS, DMSP, GAW, SS; Visualization—RFSG; Supervision—GAW, SS; Project administration—GAW, SS; Funding acquisition—GAW, SS.

## Notes

### Competing Interest Statement

The authors, Rui FS Goncalves, Raquel T. de Sousa, Daniel Stabler, Geraldine A. Wright, and Sharoni Shafir, are shareholders in Apix Biosciences, a bee-nutrition university spin-off registered in Belgium. The authors declare that they have no other competing interests. David MS Pinto has no competing interests to declare. All authors have read and approved the manuscript for submission.

### Summary of Updates

We addressed the concerns raised by the reviewers by clarifying some methodological details, such as disease management procedures and the limitations of our semi-field system. We added explanations of how Varroa load was visually assessed and kept at zero through routine monitoring and treatments, and clarified that nurse bees were sourced only from healthy, virus-free colonies. We specified that all queens were sister queens, described in detail how they were introduced and released, and clarified that all colonies began the experiment with a fully built comb. To standardise colony size, we detailed our volumetric method of measuring nurse bees (700 mL, approx. 1,580 workers per mini-nuc). We expanded the Discussion to address limitations associated with small colony size, behavioural flexibility, variation in nurse nutritional stores, mortality in enclosures, and restricted foraging. Finally, due to figure-number limits, step-by-step visual guidance on establishing mini-nucs has been moved to the supplementary materials. These revisions improve clarity, reproducibility, and transparency of the method.

