## Supplemental Figures S1, S2, S3 for "1/ A technical semi-field methodology to measure the effect of nutrition on honeybee brood rearing"

**SUPPLEMENTARY:**

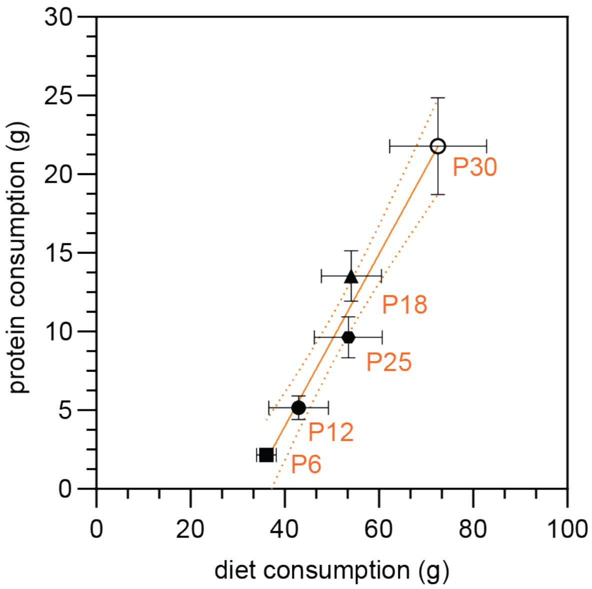


**Fig. S1. Protein-rich diets lead to greater diet consumption and protein intake.** Data points show the average results for each of the five diets varying in protein content (% w/w) inside the glasshouse: P6 (6%), P12 (12%), P18 (18%), P25 (25%), and P30 (30%) - over the 15-day period (one bioassay cycle). Orange lines depict the regression lines with 95% degree confidence between different diet types. Error bars in SEM, *n=5* mini-colonies per (P6), (P12), (P30) diet; and n=6 mini-colonies (P18), (P25) diet. The relationship between total diet consumption (x-axis) and protein intake (y-axis), where a positive association was observed - mini-colonies consumed more food and ingested significantly more protein when offered protein-rich diets.

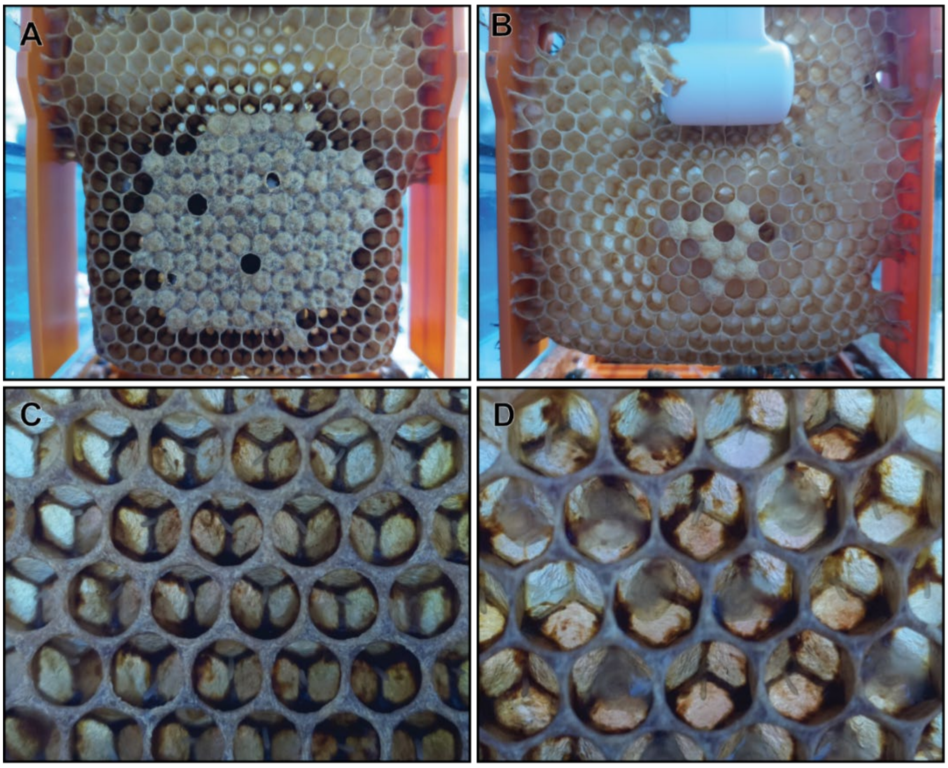


**Fig. S2. Sample pictures taken from the photographic-acrylic box device for imaging mini-colony frames.** (A) Original size picture showing a mini-colony frame with mostly capped brood and one uncapped, white-eyed pupae. (B) Original size picture showing a mini-colony frame with a white bee sensor on top, nine capped cells and several uncapped larvae. (C) Due to the concave shape of single comb cells, the zoomed-in central section picture on the mini-colony frame can display eggs, in ≈100 cells, to distinguish between empty and egg cell load . (D) zoomed-in central section picture on mini-colony frame demonstrating eggs and 1-2-day-old larvae, facility to distinguish cell load brood development in ≈100 cells.

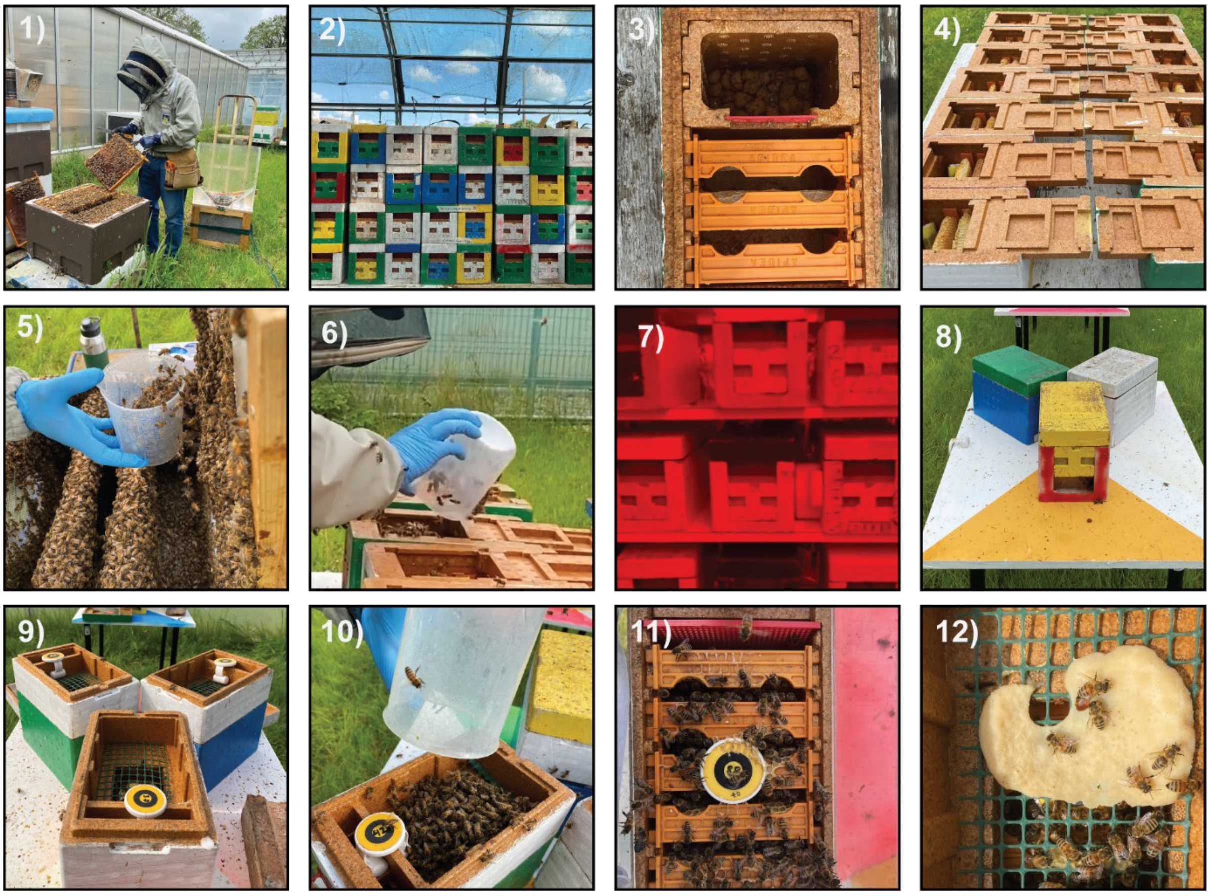

**Fig. S3. Step-by-step workflow for establishing mini-nucs inside the enclosure in the semi-field mini-colony method.**

(1) *Left:* Selection of strong, healthy colonies, with no signs of *Varroa destructor* and Deformed Wing Virus; *Right:* custom-built nurse bee collection box used for sampling.
(2) Stack of Apidea mini-nucs with different colours serving as homing cues, prepared to host nurse bees and mated queens.
(3) Top view of a mini-nuc showing clay pebbles inside the syrup feeder to prevent bee drowning (acclimation stage).
(4) Rows of Apidea mini-nucs opened bottom-up, with yellow mated-queen cages ready to be filled with nurse bees.
(5) Detail of a 500 mL beaker used to scoop bees from the nurse collection box gently.
(6) Filling each mini-nuc with a pooled mix of healthy nurse bees inside the brood chamber.
(7) Mini-nucs containing nurse bees and a mated queen inside the cold, dark room to promote nestmate recognition and colony formation.
(8) Release of mini-nucs inside the semi-field enclosure  (acclimation stage).
(9) Expansion of mini-nucs by adding two additional mini-frames and a protein patty top-feeder.
(10) Addition of nurse bees to the mini-nuc via the top feeder.
(11) Brood nest development within mini-nucs containing five comb frames, nurse bees, and a laying mated queen.
(12) Protein patty partially consumed by bees on day 7.
